# Integrative Analysis of Axolotl Gene Expression Data From Regenerative and Wound Healing Limb Tissues

**DOI:** 10.1101/693523

**Authors:** Mustafa Sibai, Cüneyd Parlayan, Pelin Tuğlu, Gürkan Öztürk, Turan Demircan

## Abstract

Axolotl (*Ambystoma mexicanum*) is a urodele amphibian endowed with remarkable regenerative capacities manifested in scarless wound healing and full restoration of amputated limbs. Several regenerative cues of the axolotl limb were successfully unraveled due to the advent of high-throughput technologies and their employment in tackling research questions on several OMICS levels. The field of regenerative biology and medicine has therefore utilized the axolotl as a major and powerful experimental model. Studies which have previously unraveled differentially expressed (DE) genes *en masse* in different phases of the axolotl limb regeneration have primarily used microarrays and RNA-Seq technologies. However, as different labs are conducting such experiments, sufficient consistency may be lacking due to statistical limitations arising from limited number of sample replicates as well as possible differences in study designs. This study, therefore, aims to bridge such gaps by performing an integrative analysis of publicly available microarray and RNA-Seq data from axolotl limb samples having comparable study designs. Three biological groups were conceived for the analysis; homeostatic tissues (control group), from amputation/injury timepoint up to around 50 hours post amputation (wound healing group), and from 50 hours to 28 days post amputation/injury (regenerative group). Integrative analysis was separately carried out on the selected microarray and RNA-Seq data from axolotl limb samples using the “merging” method. Differential expression analysis was separately implemented on the processed data from both technologies using the R/Bioconductor “limma” package. A total of 1254 genes (adjusted P < 0.01) were found DE in regenerative samples compared to the control, out of which 351 showed magnitudes of Log Fold Changes (LogFC) > 1 and were identified as the top DE genes from data of both technologies. Downstream analyses illustrated consistent correlations of the logFCs of DE genes distributed among the biological comparisons, within and between both technologies. Gene ontology annotations demonstrated concordance with the literature on the biological process involved in the axolotl limb regeneration. qPCR analysis validated the observed gene expression level differences between regenerative and control samples for a set of five genes. Future studies may benefit from the utilized concept and approach for enhanced statistical power and robust discovery of biomarkers of regeneration.

## 1. INTRODUCTION

*Ambystoma mexicanum* (Axolotl) is a salamander species of amphibians which has recently been established as a promising vertebrate model system for developmental and regenerative biology due to its unique features of high regenerative capacity, scarless wound healing, and low cancer incidence [1,2]. What distinguishes axolotls is the preservation of their juvenile features when they reach sexual maturity [3]. Due to their inability to undergo natural metamorphosis, axolotls keep on exhibiting embryonic-like cell characteristics, which promotes the finely-tuned regenerative capacity of their body parts throughout their lifespan [1,4]. Organs such as the heart [5–7], spinal cord [8], and even brain [9] are prime examples of what axolotls can faithfully regenerate besides their limbs. It’s been suggested that, unlike the amniote vertebrate, the successive regenerative capacity of axolotls may be driven by a weak inflammatory response due to their simpler adaptive immune system [1,10]. In response to experimental induction of metamorphosis via thyroid hormone administration, diminished regenerative power of axolotls is observed for some body parts such as appendages [11,12], while such regenerative potential seems to be unobstructed for other parts of the body [1,13].

The process of axolotl limb wound healing and regeneration involved several key genes [14–20] and proceeds in several stages. In response to an amputation, a thin wound epidermis forms around the severed stump within 24 hours due to the migration of a collection of epidermal cells to the amputation site. The newly constructed epidermis substantially differs structurally and molecularly from an intact, fully differentiated epidermis [21,22]. Within 48 hours thereafter, the wound area is infiltrated by macrophages where they phagocyte the debris of dead cells and clear the wound zone from different kinds of pathogens. Such an indispensable role of macrophages for successful regeneration has been demonstrated by previous studies [23]. In the following couple of days, several key processes such as activation of progenitor cells as well as dedifferentiation of terminally differentiated cells take place as a result of secretion of mitogens and growth factors from the epidermis accompanied by innervation [22]. These processes lead to the formation of what’s known as the blastema, which is characterized by highly proliferative, heterogenous cells that are morphologically similar to fibroblasts [22,24]. They are encoded with precise positional information, behave as autonomous units, and exhibit unidirectional signaling driven by factors originating from wound epidermis [22]. After blastema cells reach a definitive size, they flatten out for cartilage to condense, allowing the differentiation of the required tissue types, and ultimately constructing a delicately-sized, perfectly regenerated limb that is identical to the amputated one [22].

Axolotls can be easily bred year-round and their generation time is not more than a year [22,25]. Multiple studies have been successful in applying transgenesis and genome editing techniques on these animals [26–30]. While the genome of the axolotl is a simple diploid with 14 pairs of chromosomes, its enormous, highly repetitive sequence and long introns [31] have been the major hurdles towards obtaining a full genome assembly due to the inability of acquiring sufficient read-length and the absence of an improved methodology of genome assembly [22]. Consequently, only very recently sequencing and assembly of axolotl genome was reported [32]. Therefore, proteomics, transcriptomics (RNA-Seq), and microarrays have been the primary, indispensable tools for investigating gene networks and pathways of axolotl’s regenerative mechanisms [17,22,33–36]. While microarrays can be used to design probes for any condition of interest, RNA-Seq facilitates the detection of a wider range of expression values at a single-nucleotide resolution as well as accurately detects highly expressed genes with no saturation effect that is typically observed in microarrays [22]. Moreover, RNA-Seq allows for novel discoveries of mRNAs particularly for non-model systems and has the ability of quantifying isoforms of genes. Recent advances paved the way for even using individual cells as the source of low RNA input, which resulted in an illuminating understanding of the specific cells and tissues contributing to the process of limb regeneration [22,37].

Despite such advancements and success in utilizing microarrays and RNA-Seq technologies in the quest for unraveling the mechanisms of axolotl limb regeneration, a strong and popular methodology that has the potential for enhancing our current knowledge on limb regeneration is missing in the axolotl literature. Integrative data analysis (IDA) is a key methodology that is applied across many scientific disciplines and aims to derive scientific consensus on a particular research question [38–40]. Although the concept of IDA has recently been expanded to refer to experiments aiming to integrate information from several layers of “omics” information (aka multi-omics) [41], the utilized IDA in this study refers to the process of combining information from different platforms across independent studies [40]. The latter IDA concept is commonly used in biomedical sciences to detect differentially expressed (DE) genes for having a better gene signature for basic science and clinical applications [40,42,43]. As microarray and RNA-Seq data are increasingly becoming publicly available, IDA can benefit the axolotl regeneration field by improving candidate gene selection and establishing biomarkers for different stages of limb regeneration.

Factors such as directly-incomparable experimental techniques between different studies due to poorly articulated experimental designs as well as data deposition in public databases with no relevant publications are the main reasons why many biological research and preclinical medical sciences have not embraced the application of IDA very quickly [38]. Therefore, when such factors are no longer an obstacle, the advantages of implementing IDA can be realized from the limitations of individual studies. First and foremost, the cost of utilizing new technologies often times leads to the collection of a limited number of replicates [38]. Moreover, researchers are faced with challenging statistical issues arising from such limited replicates, manifested in high false-positive and false-negative observations [38,44]. Lastly but not least, an overlap of experimental design of individual studies may in some cases generate similar data, due to lack of an established IDA pipeline. Therefore, when IDA is applied, sample size increases, individual study-specific biases are minimized, and more statistical power is achieved [39,40,45–47]. Meta-analysis and cross-platform normalization (aka “merging”) are two fundamental approaches to perform IDA [40]. While meta-analysis integrates statistics from different studies at a “late stage”, merging integrates data before running the statistical test [40]. It has been argued that whenever datasets are selected for answering particularized questions and are reasonably homogenous, the merging method outperforms meta-analysis for biomarker discovery analyses [40,48].

In this study, we aimed to identify the genes that could be considered as biomarkers for the regenerative phase of axolotl limb which starts around 2 dpa until 28 dpa by separately implementing integrative data analysis on publicly available microarray and RNA-Seq axolotl data using cross-platform normalization (merging) methodology. We report the finding of 1254 differentially expressed (DE) genes (adjusted P < 0.01), commonly identified by microarray and RNA-Seq integrative data analysis in the regenerative phase compared to the control (intact) limb. In addition, out of those 1254 DE genes, the top 351 genes enriched in several biological processes (adjusted p-value < 0.05) such as regulation of cell cycle processes, mitotic nuclear division, regulation of mRNA metabolic processes, muscle filament sliding and contraction, muscle structure development, and cardiac muscle tissue morphogenesis. Overall, this report identifies a set of commonly found differentially expressed genes curated from various studies. Our methodological approach would be benefited by the regeneration community, offering a closer look at the differentially expressed genes that are present in up-to-date high-throughput gene expression studies.

## 2. METHODS

### 2.1 Gene-expression Data Collection

Gene Expression Omnibus (GEO) database [49,50] (http://www.ncbi.nlm.nih.gov/geo/) and the European Nucleotide Archive (ENA) database [51] (https://www.ebi.ac.uk/ena) were used to collect microarray and RNA-Seq axolotl data, respectively. The collected data were subjected to selective criteria filtering which was set according to *Preferred Reporting Items for Systematic Reviews and Meta-Analyses (PRISMA)* [52], based on which GEO series were collected as follows:

1. GEO series data deposited until September 2018.
2. Non-redundant series.
3. Series pertinent to Axolotl tissues.
4. Series having unduplicated datasets.

The number of GSM files representing our meta-data design (group types), how many of each group belong to which sub-platforms and to which study (series), is provided in **Table 1** for Microarray technology [16,53–55] and in **Table 2** for RNA-Seq technology [17,36,53,56,57]. The accession number of the GEO series (GSE number), title, publication, number of the selected samples, number of the total samples, platform type, link to the gene-expression data, and the year of each study can be found in **Supplementary Table 1** for Microarray technology and in **Supplementary Table 2** for RNA-Seq technology. Further detailed information such as GSM accessions and timepoints are provided in **Supplementary Tables 3 and 4** for Microarray and RNA-Seq technologies, respectively.

**Table 1:**
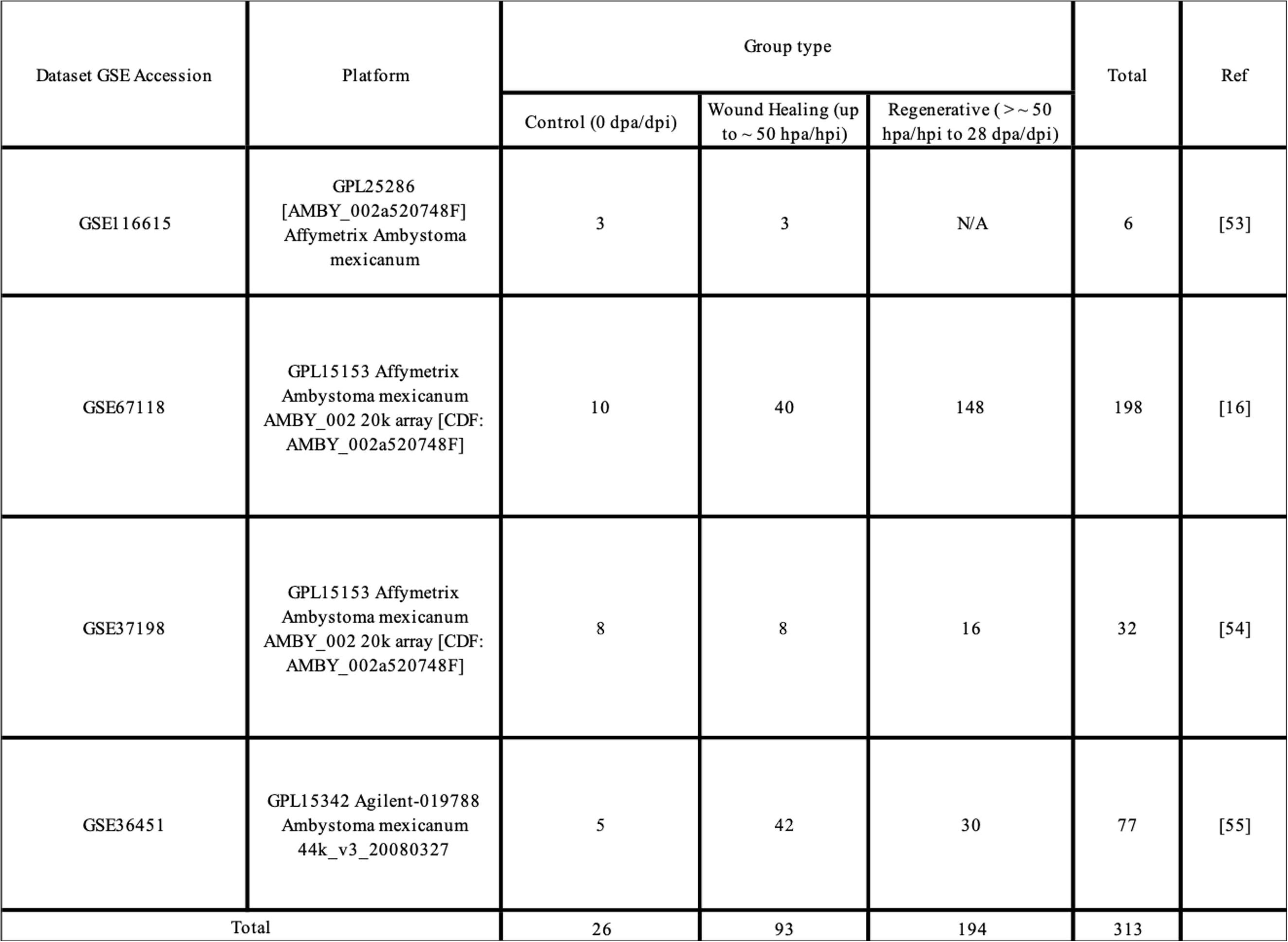
Summary of the selected microarray data from axolotl samples used for integrative analysis

**Table 2:**
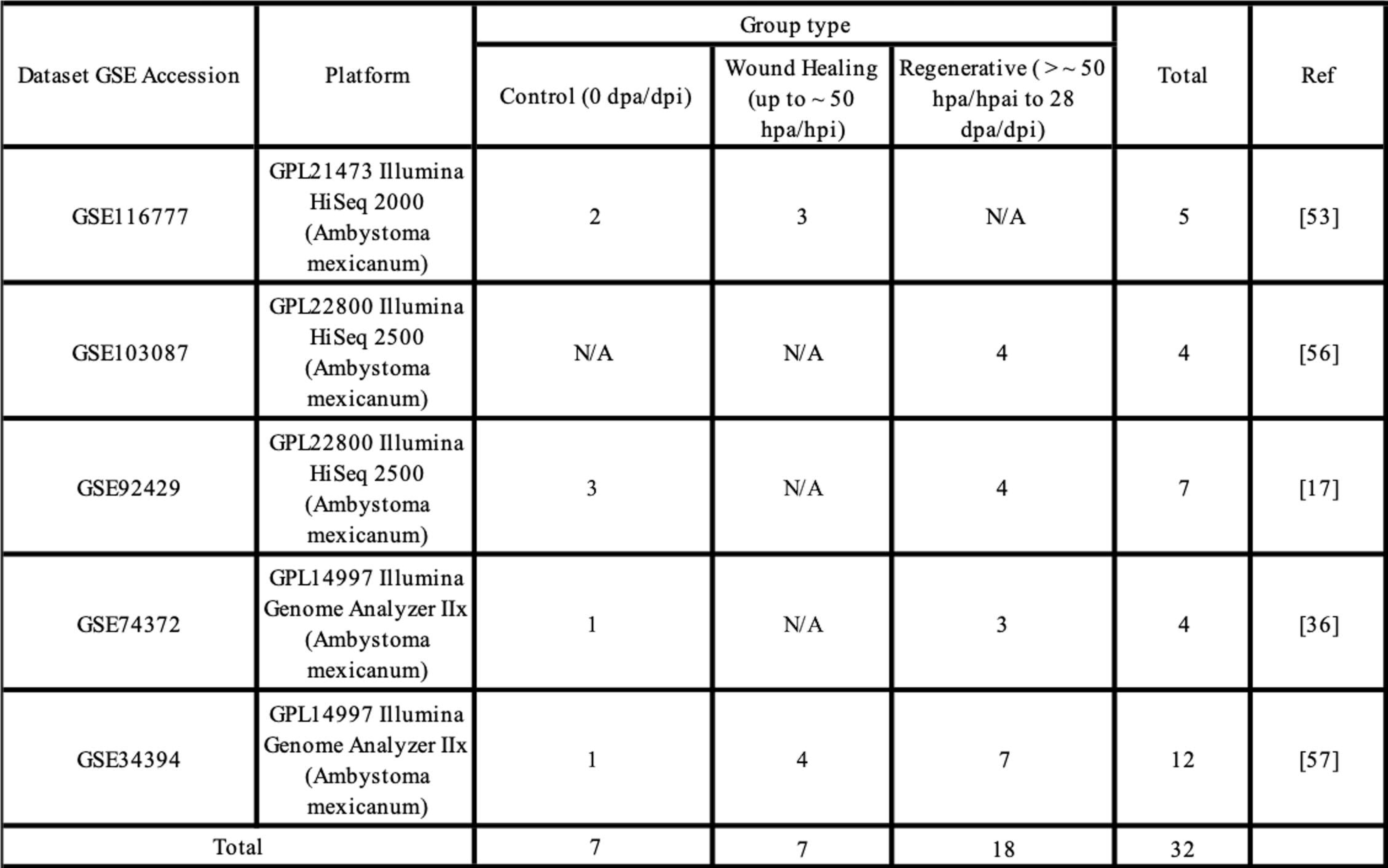
Summary of the selected RNA-seq data from axolotl samples used for integrative analysis

### 2.2 Gene-expression Data Processing

R programming environment (version: 3.5.1) [58], packages from open development software Bioconductor Project [59,60], and parts of a published gene-expression analysis workflow [61] were used for integrative data analyses of microarray and RNA-Seq axolotl data.

The full axolotl microarray data processing workflow can be found in **Supplementary Figure 1**. CEL files from the 3 Affymetrix datasets (GSE116615, GSE67118, GSE37198) were processed as a single dataset having a total of 236 samples (21 control, 51 wound healing, 164 regenerative) and 20080 probesets. Summarization, quantile-normalization, and log2-transformation were applied on this dataset by using the Robust Multichip Average (RMA) algorithm from the Bioconductor’s “affy” package in R [62]. Samples from the Agilent dataset (GSE36451) were collected from GEO as summarized, quantile-normalized, and log2-transformed, having a total of 77 samples (5 control, 42 wound healing, 30 regenerative) and 43796 probesets. The resultant two datasets (Affymetrix and Agilent) were filtered for lowly-expressed probesets using Bioconductor’s “Biobase” package in R [59]. Probesets with intensities higher than a manual threshold of a “4” median intensity in at least as many samples of the smallest group were kept, resulting in 17658 probesets for Affymetrix dataset and 41579 probesets for Agilent dataset. The former probesets were annotated using AMBY_002a520748F Affymetrix probeset annotation file (∼ 20k probesets) provided by Sal-Site (http://www.ambystoma.org/genome-resources/20-gene-expression) and the latter probesets were annotated using GPL15342 Agilent annotation file obtained from GEO (https://www.ncbi.nlm.nih.gov/geo/query/acc.cgi?acc=GPL15342). The annotation yielded 13316 Affymetrix and 21419 Agilent probeset-gene mappings including duplicates, respectively. The “WGCNA” R package [63] was used to remove those duplicates by respectively collapsing the Affymetrix and Agilent data from probeset-level to gene-level while taking the maximum row mean value of the duplicated probesets to represent the corresponding gene, yielding a total of 10442 Affymetrix and 5083 Agilent unique genes. The latter two gene lists were then merged together resulting in 4322 unique genes common between Affymetrix and Agilent datasets. Thereafter, the gene lists of both of the Affymetrix and Agilent log2-transformed data were each substituted with the 4322 common gene list, followed by transforming each of them to z-scores [64] in order to minimize the batch effect between the two platforms. The resultant two z-scored Affymetrix and Agilent lists were merged together, yielding a single dataset of 313 samples (26 control, 93 wound healing, 194 regenerative) and 4322 genes.

**Figure 1:**
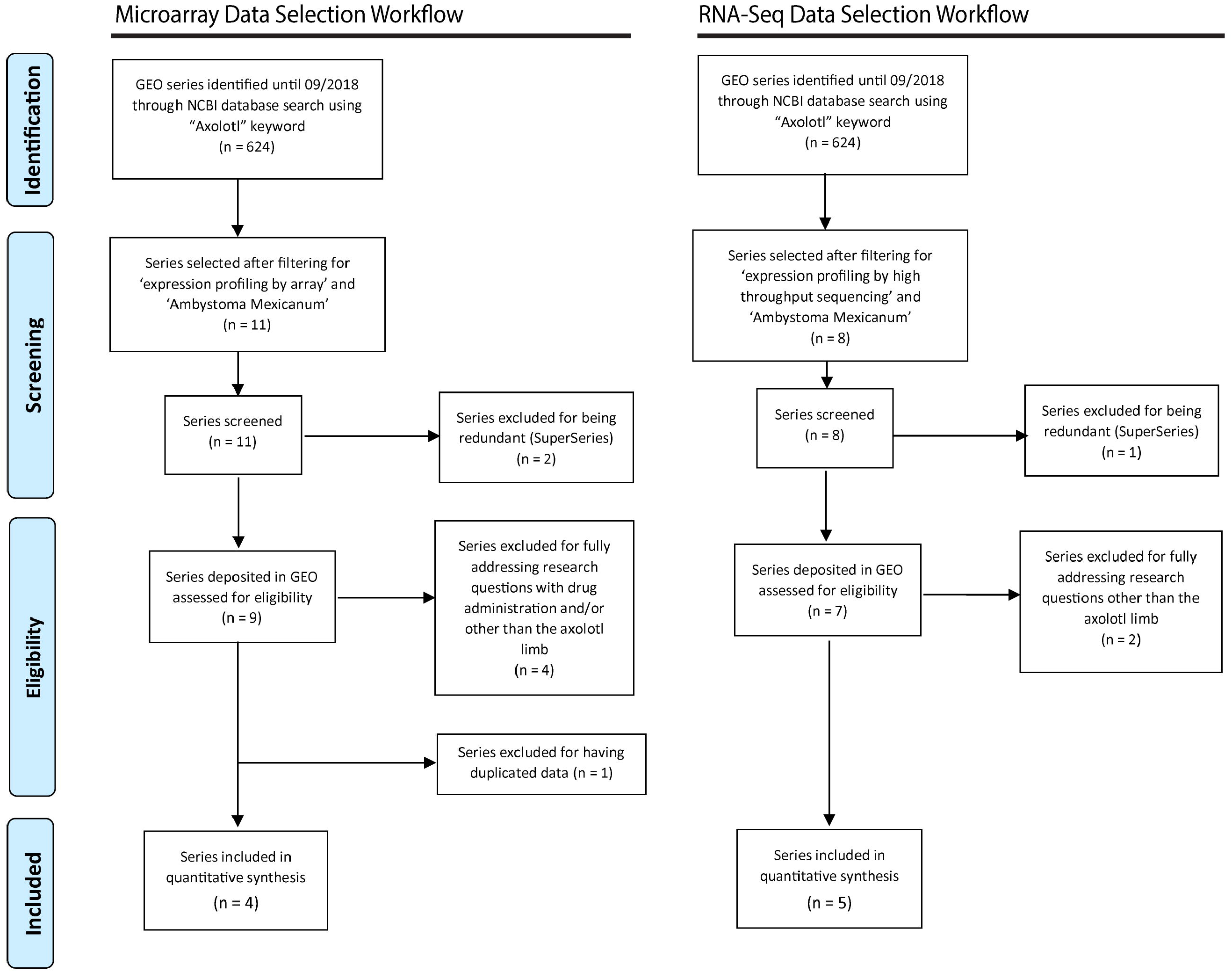
Flow diagram illustrating the selection criteria used for separately performing integrative analysis on microarray and RNA-seq data from axolotl samples. The diagram is prepared according to *Preferred Reporting Items for Systematic Reviews and Meta-Analyses (PRISMA)*

The steps of axolotl RNA-Seq data processing workflow can be found in **Supplementary Figure 2**. Fastq files corresponding to samples from the 5 GSE datasets were obtained from the European Nucleotide Archive (ENA) database. Datasets with paired-end libraries are GSE116777 (2 control, 3 wound-healing), GSE92429 (3 control, 4 regenerative), and GSE103087 (4 regenerative), whereas both GSE74372 (1 control, 3 regenerative) and GSE34394 (1 control, 4 wound-healing, 7 regenerative) are single-end libraries. Recently, an axolotl transcriptome “V5 contig assembly” with contig length of 19,732 bp was published by Dwaraka *et al*., [53] and was chosen to be our transcriptome reference. According to the authors, a total of 31,886 pairwise alignments with more than 98 % sequence similarity were detected between V5 RNA-Seq contigs and the 20,036 V3 contigs which were used to design Affymetrix microarray probesets (AMBY_002a520748F) already having an annotation file. Therefore, since our microarray analysis pipeline included those microarray probesets along with its annotation file, a new annotation file for the 31,886 aligned contigs of the V5 assembly was generated for our RNA-Seq analysis pipeline by merging the 31,886 V5 contigs-V3 probesets alignment with V3 (AMBY_002a520748F) annotation file, resulting in around 25,000 genes. The transcriptome-wide quantifier “Salmon” tool [65] was then used to separately quantify the expression of the transcripts of each of the 5 datasets by indexing the V5 transcriptome upon which direct quantification of the reads was carried out. The R/Bioconductor “tximport” package [66] was used to import the scaled counts generated from transcript abundances along with our RNA-Seq annotation file, yielding a count matrix of all samples from the 5 datasests combined (32 samples) with 10,000 unique genes. Lowly expressed genes were filtered out by keeping genes having minimum counts for at least some samples using R/Bioconductor’s “edgeR” package [67,68]. The filtered count data was then converted to log2-counts per million (logCPM) and was made ready to be used for linear modeling for differential expression analysis using the “voom” function from R/Bioconductor’s “limma” package [69–71].

**Figure 2:**
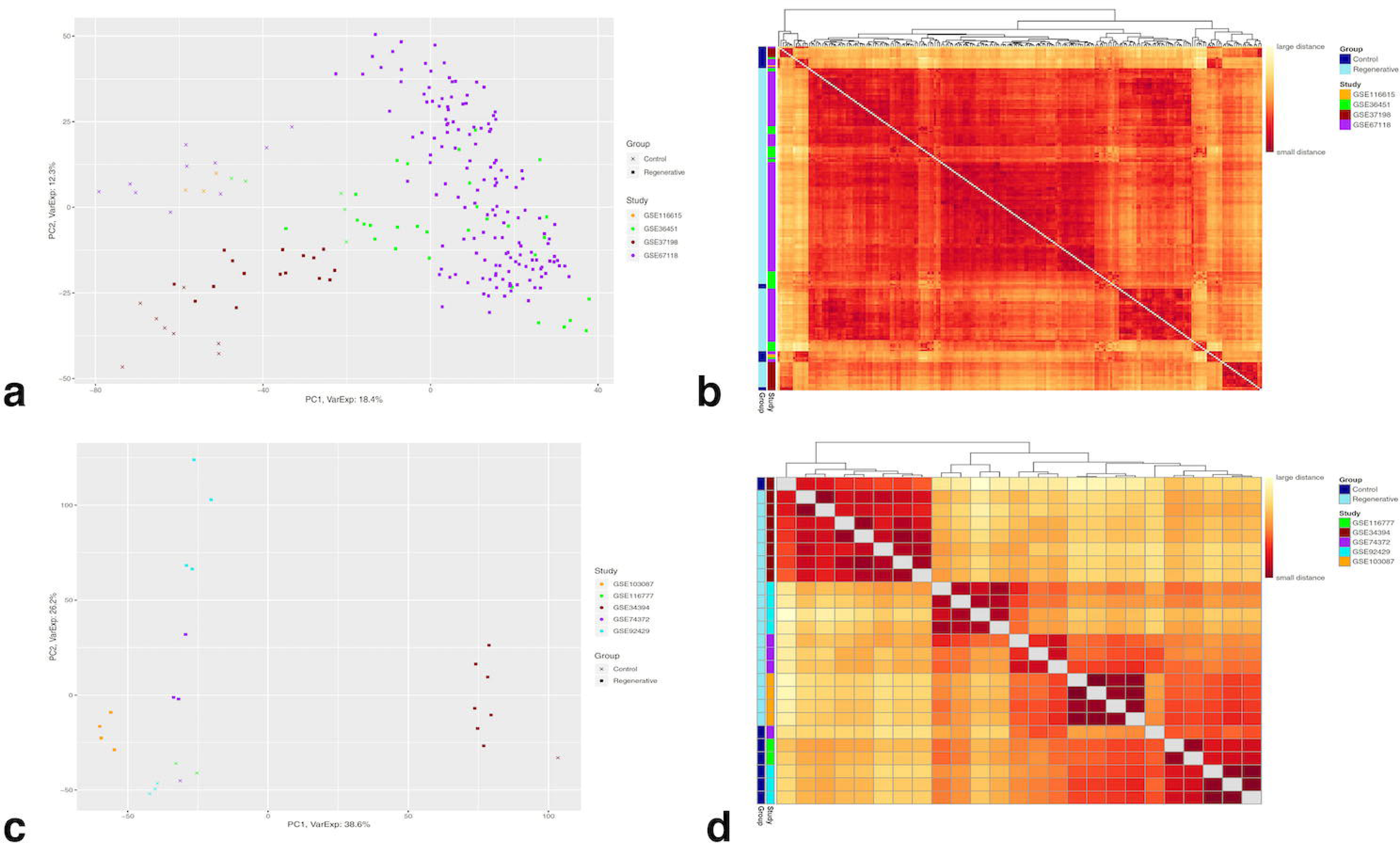
Whole gene expression data-based principal component analysis and sample-to-sample clustering heatmaps for control and regenerative samples of A,B) Microarray quantile-normalized, log2-transformed, z-scored, data (4322 genes, 220 samples) and of C,D) RNA-seq logCPM (voom) counts (7562 genes, 25 samples). A,C) principle component analysis. B,D) sample-to-sample clustering heatmap. The number of samples per group are; 26 control and 194 regenerative for microarray data; 7 control and 18 regenerative for RNA-Seq data.

### 2.3 Differential Expression Analysis (DEA)

Microarray DEA was performed on the z-scored, combined data (313 samples, 4322 genes), while RNA-seq DEA was performed on the logCPM (voom) counts (32 samples, 7562 genes), respectively. The following analyses were performed using the R/Bioconductor “limma” package [69–71]. Three biological group comparisons were conceived for DEA of both microarray and RNA-Seq data. First of which aims to observe gene expression changes in wound healing tissues compared to homeostatic tissues (control group). The second comparison aims to observe gene expression changes in regenerative tissues group compared to control group. The third comparison aims to observe gene expression changes occurring in regenerative tissues compared to wound healing tissues. Design matrices were incorporated for each contrast to account for group type and study origin (batch factor) of every sample, followed by a fitted linear model on the expression data for each gene, which were then ranked based on an order of evident differential expression by applying the empirical Bayes method. False discovery rate (FDR) using the Benjamini-Hochberg (BH) method was controlled at 0.01, below which all genes were extracted representing lists of DE genes for each comparison from microarray and RNA-Seq data.

### 2.4 Principal Component Analyses, Clustered Heatmaps, and Correlations

In order to explore overall relationship occurring among samples of the z-scored microarray data and that of the “voom” counts RNA-Seq data, for each comparison, principal component analysis (PCA) was separately implemented on each of them in base R. PCA was also separately used on DE microarray and DE RNA-Seq data while minimizing the “study origin” batch effect using “limma” R package [69] to illustrate how differentially expressed genes determine the clustering among samples, for every comparison. Complementary to both PCA approaches, heatmaps with Sample-to-Sample clustering based on “Manhattan” distance were implemented using “pheatmap” R package [72]. The distribution and correlation of the DE genes among the three comparisons within each technology and the subsequent comparison of correlation of the DE genes between the two technologies were carried out using Venn diagram online tool (http://bioinformatics.psb.ugent.be/webtools/Venn/) and scatter plots using the R/Bioconductor “ggplot2” package [73]. Pair-wise correlation testing of logFC of the DE genes for the three comparisons within and between the two technologies was performed using the “Spearman” correlation test in base R.

### 2.5 Gene Ontology Enrichment Analysis

Gene Ontology annotations of the top DE genes commonly identified by microarray and RNA-Seq DE analyses were carried out using Bioconductor’s “clusterProfiler” package in R [74]. The top up-regulated and down-regulated Entrez ID-mapped genes of each comparison were separately queried against the three GO categories (biological process “BP”, cellular component “CC”, and molecular function “MF”). The background gene list was all Entrez IDs from axolotl Affymetrix (AMBY_002a520748F) annotation file. The organism database was set as homo sapiens “org.Hs.eg.db”, the adjusted p value method was Benjamini & Hochberg (BH), and cutoffs for p and q values were set to 0.05. Redundant GO terms were eliminated using “simplify” function from the clusterProfiler package [74]. The latter package was also used to visualize some GO categories and genes using a heatmap-like plot (heatplot) and a circular net (cnetplot). Bar plots of the top 10 terms of GO categories were generated using the R/Bioconductor “ggplot2” package [73].

### 2.6 Heatmap Generation for Top 100 Regenerative vs. Control Genes

The top 100 DE genes detected by DEA of each technology in regenerative vs. control comparison were visualized in a clustered heatmap using “pheatmap” package in R [72]. The samples were clustered using the “Manhattan” distance measure. The genes were hierarchically clustered using “Complete Linkage” method. The values were centered and scaled in the row direction.

### 2.7 qPCR analysis

RNA was isolated from axolotl limb tissues (0,1 and 7 dpa) using TRIzol reagent (Invitrogen). RNA quantity was checked by spectrophotometry using a NanoDrop ND-1000 (NanoDrop) and quality of isolated RNA was assessed by gel electrophoresis. M-MLV Reverse Transcriptase (Thermo Fisher Scientific) was used to perform reverse transcription according to manufacturer’s procedure. Quantitative PCR assays were performed at following conditions: initial denaturation at 95 °C for 2 minutes, and 40 cycles of denaturation at 95 °C for 5 seconds, annealing at 55 °C for 10 seconds and extension at 72 °C for 15 seconds. For qPCR reactions, SensiFAST™ SYBR® No-ROX Kit (BIOLINE, BIO-98005) and CFX Connect Real-Time PCR Detection System (BIO-RAD) was used. Gene expression levels were calculated using the 2–ΔΔCt method and cDNA concentrations were normalized with Ef1-α (elongation factor 1-alpha) housekeeping gene. Primers used in this study are listed in **Supplementary Table 5**.

## 3. RESULTS

### 3.1 Gene-expression Data Selection and Search Criteria

Three biological groups were set for our integrative data analysis; “control” group (intact/amputated/injured limbs and/or flank wounds at 0-hour timepoints, “wound healing” group (amputated/injured limbs and/or flank wounds up to around 50-hours post amputation/injury, and “regenerative” group (amputated/injured limbs and/or flank wounds of time points ranging from around 50 hours to 28 days post amputations/injuries). According to the criteria for microarray data selection (**Figure 1**), a total of 4 GSE datasets (series) were selected, three of which were based on the Affymetrix Ambystoma mexicanum platform of slightly different versions; GSE116615 is based on GPL25286 [AMBY_002a520748F], while GSE67118 and GSE37198 are both based on GPL15153 [AMBY_002a520748F]. The fourth study (GSE36452) is based on the Agilent-019788 Ambystoma mexicanum platform of GPL15342 [44k_v3_20080327] version. The excluded samples out of the selected microarray datasets were those of denervated limbs (in GSE37198) which can’t regenerate and those of limb buds (in GSE36451) that are totally distinct from a fully mature, amputated limb. Overall, 6 samples from GSE116615 dataset, 198 samples from GSE67118 dataset, 32 samples from GSE37198 dataset, and 77 samples from GSE36451 dataset were selected, forming a total of 313 samples. Out of those, 26 are control samples, 93 are wound healing samples, and 194 are regenerative samples, all of which to be used for the integrative microarray analysis (**Table 1**).

As for the criteria for RNA-Seq data selection (**Figure 1**), a total of 5 GSE datasets (series) were selected, all of which were based on slightly different versions of the Illumina *Ambystoma mexicanum* platform; GSE116777 used GPL21473 HiSeq 2000, GSE103087 and GSE92429 both used GPL22800 HiSeq 2500, and GSE74372 and GSE34394 both used GPL14997 Genome Analyzer IIx. The excluded samples out of the selected RNA-Seq datasets were those identified as an outlier (in GSE116777) by the authors [53], those which underwent several rounds of amputation-regeneration (in GSE103087), those which were prepared for small RNA (sRNA) profiling experiments (in GSE74372), those which were mouse samples (in GSE34394), and those extracted from multiple positions along the axolotl limb except for those of the upper-arm to be used in the control group as well as the proximal and distal blastemas to be used in the regenerative groups (in GSE92429). Overall, 5 samples from GSE116777 dataset, 4 samples from GSE103087 dataset, 7 samples from GSE92429 dataset, 4 samples from GSE74372 dataset, and 12 samples from GSE34394 dataset were selected, forming a total of 32 samples. Out of those, 7 are control samples, 7 are wound healing samples, and 18 are regenerative samples, all of which to be used for the integrative RNA-Seq analysis (**Table 2**).

Further detailed information about the selected microarray and RNA-Seq studies on the overall design, selected and total number of samples, and the biological group types we conceived for the integrative analysis are summarized in **Supplementary Tables 1-4**.

### 3.2 Whole Gene Expression Data-based PCA and Heatmap Clustering

The first principal component (PC1) as well as the sample-to-sample clustering heatmap of the z-scored, microarray data (4322 genes) show a rough separation among the samples based on their group types; for control and regenerative samples (**Figure 2A,B**) and for the other two group pairs (control and wound healing, regenerative and wound healing) (**Supplementary Figures 3A,B and 4A,B**). However, the second principal component (PC2) seems to separate the samples based on their study origin. Therefore, the source of variation is likely explained by differential gene expression between the group types.

**Figure 3:**
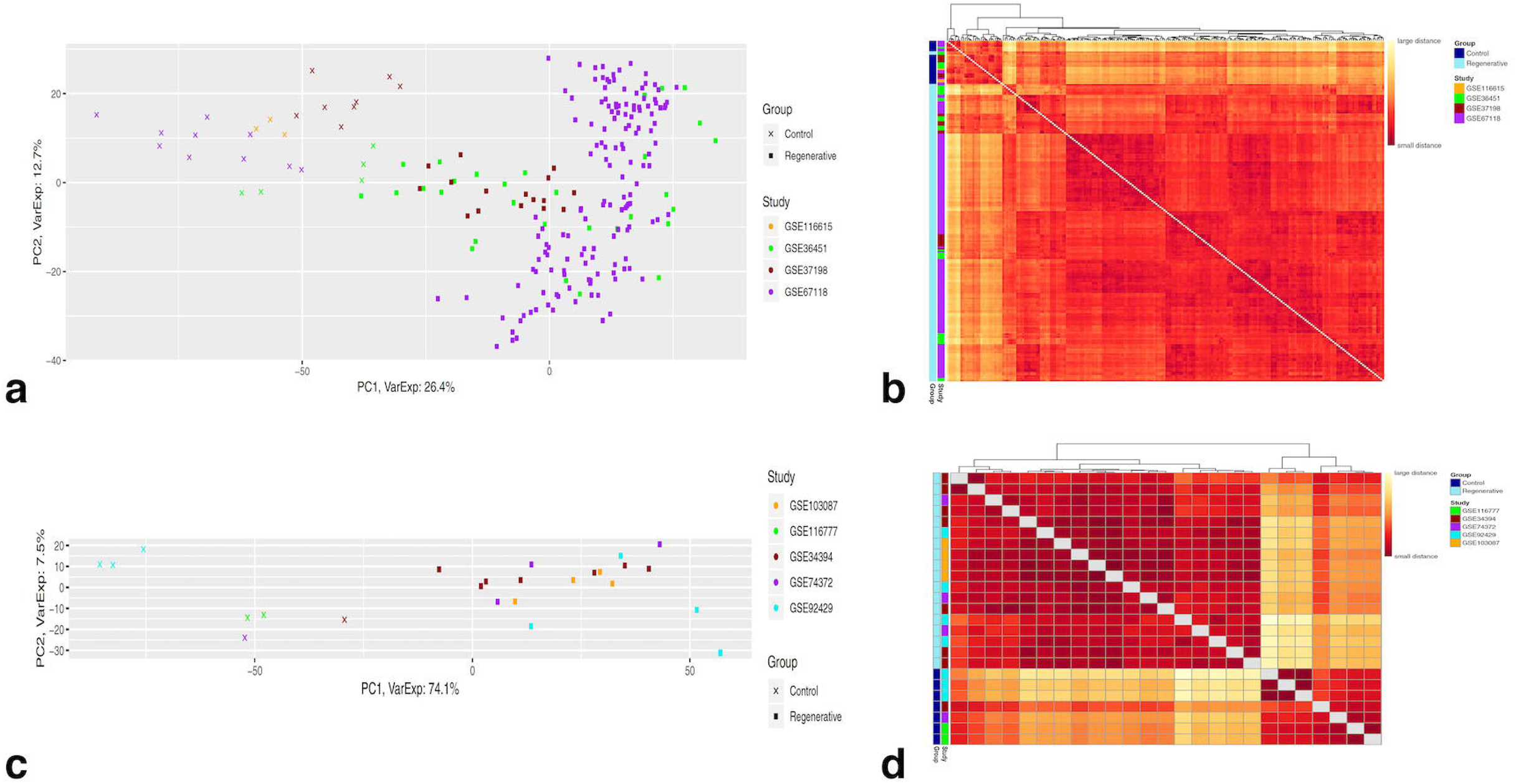
DEA-based principal component analysis and sample-to-sample clustering heatmaps for regenerative vs. control comparison of A,B) Microarray quantile-normalized, log2-transformed, z-scored data after DEA (2748 DE genes, 220 samples) and of C,D) RNA-seq logCPM (voom) counts data after DEA (2992 DE genes, 25 samples). A,C) principle component analysis. B,D) sample-to-sample clustering heatmap. The number of samples per group are; 26 control and 194 regenerative for microarray data; 7 control and 18 regenerative for RNA-Seq data. The DE genes have an adjusted p-value < 0.01.

**Figure 4:**
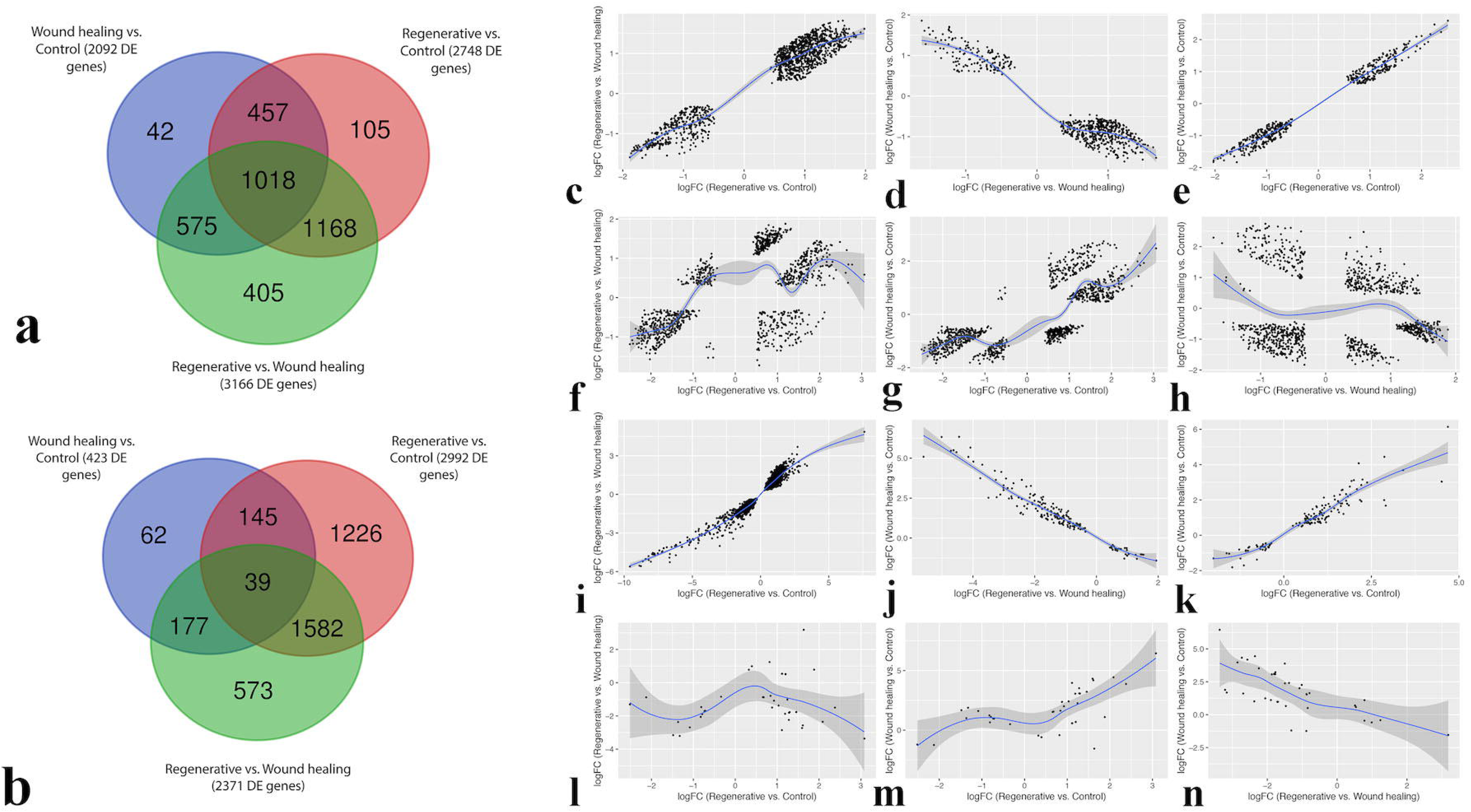
Venn Diagram of the distribution of significant genes (adj.p <0.01) among the three comparisons, from A) Microarray data and B) RNA-Seq data; A,B) Each complete circle represents the number of differentially expressed genes of a certain comparison as resulted by DEA. C-H) Scatter plots of the log-fold change of significant genes (adj.p <0.01) shared by two or more comparisons from microarray data (as shown in A). C) Scatter plot of the logFC of the 1168 DE genes shared by Regenerative vs. Control and Regenerative vs. Wound healing comparisons. D) Scatter plot of the logFC of the 575 DE genes shared by Regenerative vs. Wound healing and Wound healing vs. Control comparisons. E) Scatter plot of the logFC of the 457 DE genes shared by Regenerative vs. Control and Wound healing vs. Control comparisons. I-N) Scatter plots of the log-fold change of significant genes (adj.p <0.01) shared by two or more comparisons from RNA-Seq data (as shown in B). I) Scatter plot of the logFC of the 1582 DE genes shared by Regenerative vs. Control and Regenerative vs. Wound healing comparisons. J) Scatter plot of the logFC of the 177 DE genes shared by Regenerative vs. Wound healing and Wound-healing vs. Control comparisons. K) Scatter plot of the logFC of the 145 DE genes shared by Regenerative vs. Control and Wound healing vs. Control comparisons. F-H, L-N) Pair-wise scatter plots of the logFC of the 1018 and 39 DE genes shared by all three comparisons from microarray and RNA-Seq data, respectively; F,L) Regenerative vs. Control and Regenerative vs. Wound healing, G,M) Regenerative vs. Control and Wound healing vs. Control, H,N) Regenerative vs. Wound healing and Wound healing vs. Control.

On the other hand, the separation among the samples displayed by the first principal component (PC1) in addition to the sample-to-sample clustering heatmap of the voom-counts, RNA-Seq data (7562 genes) seems to be influenced by their study origin, conspicuously between GSE34394 and the rest; for control and regenerative samples (**Figure 2C,D**) and for the other two group pairs (control and wound healing, regenerative and wound healing) (**Supplementary Figures 3C,D and 4C,D**). The second principal component (PC2), however, seems to roughly separate the samples based on their group types.

The whole gene expression data-based PCA and clustering approach appears to emphasize an overall batch factor (study origin) which has a more dominant effect on how samples are separated than the time points (group types) in both microarray and RNA-Seq data.

### 3.3 DEA-based PCA and Heatmap Clustering

DEA of the z-scored, microarray data resulted in 2748 DE (adj p-value < 0.01) genes in regenerative vs. control, while 2092 genes and 3166 genes were DE (adj p-value < 0.01) in wound healing vs. control and regenerative vs. wound healing, respectively. Furthermore, DEA of the voom-counts, RNA-Seq data yielded 2992 DE (adj p-value < 0.01) genes in regenerative vs. control, while 423 genes and 2371 genes were DE (adj p-value < 0.01) in wound healing vs. control and regenerative vs. control, respectively. The result of DEA for every comparison showed p-value enrichment near-zero peak corresponding to the number of DE genes from microarray data (**Supplementary Figure 5**) and RNA-Seq data (**Supplementary Figure 6**), among which the aforementioned number of DE genes were considered significant (adj p-value < 0.01).

**Figure 5:**
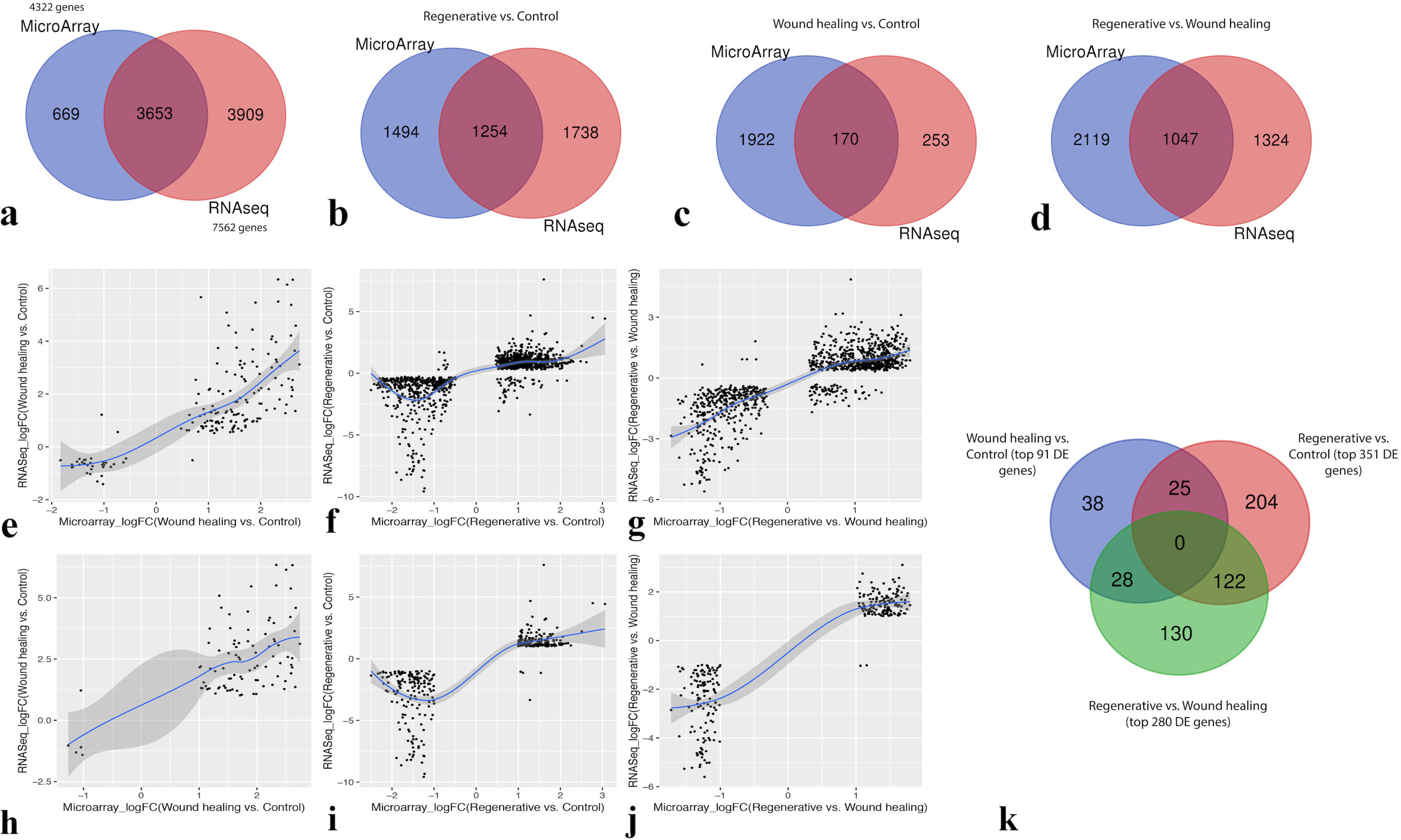
A) Venn Diagram of the initial number of genes used before differential expression analysis. This number for each platform is obtained just after the filtering and /or merging. B-D) Venn diagrams of the distribution of DE genes (adj p-value < 0.01) between Microarray and RNA-seq data as well as E-G) scatter plots of the log-fold change of the common DE genes between the two technologies for B,E) wound healing vs. control, C,F) regenerative vs. control, and D,G) regenerative vs. wound healing. H-J) Scatter plots of the log-fold change of the top DE genes (adj p-value < 0.01, logFC magnitudes > 1) commonly identified by Microarray and RNA-seq technologies’ analyses for H) wound healing vs. control (91 top DE genes), I) regenerative vs. control (351 top DE genes), and J) regenerative vs. wound healing (280 top DE genes). K) Venn Diagram of the distribution of the top DE genes (adj p-value < 0.01, logFC magnitudes > 1) commonly identified by the analyses of both technologies among the three comparisons.

**Figure 6:**
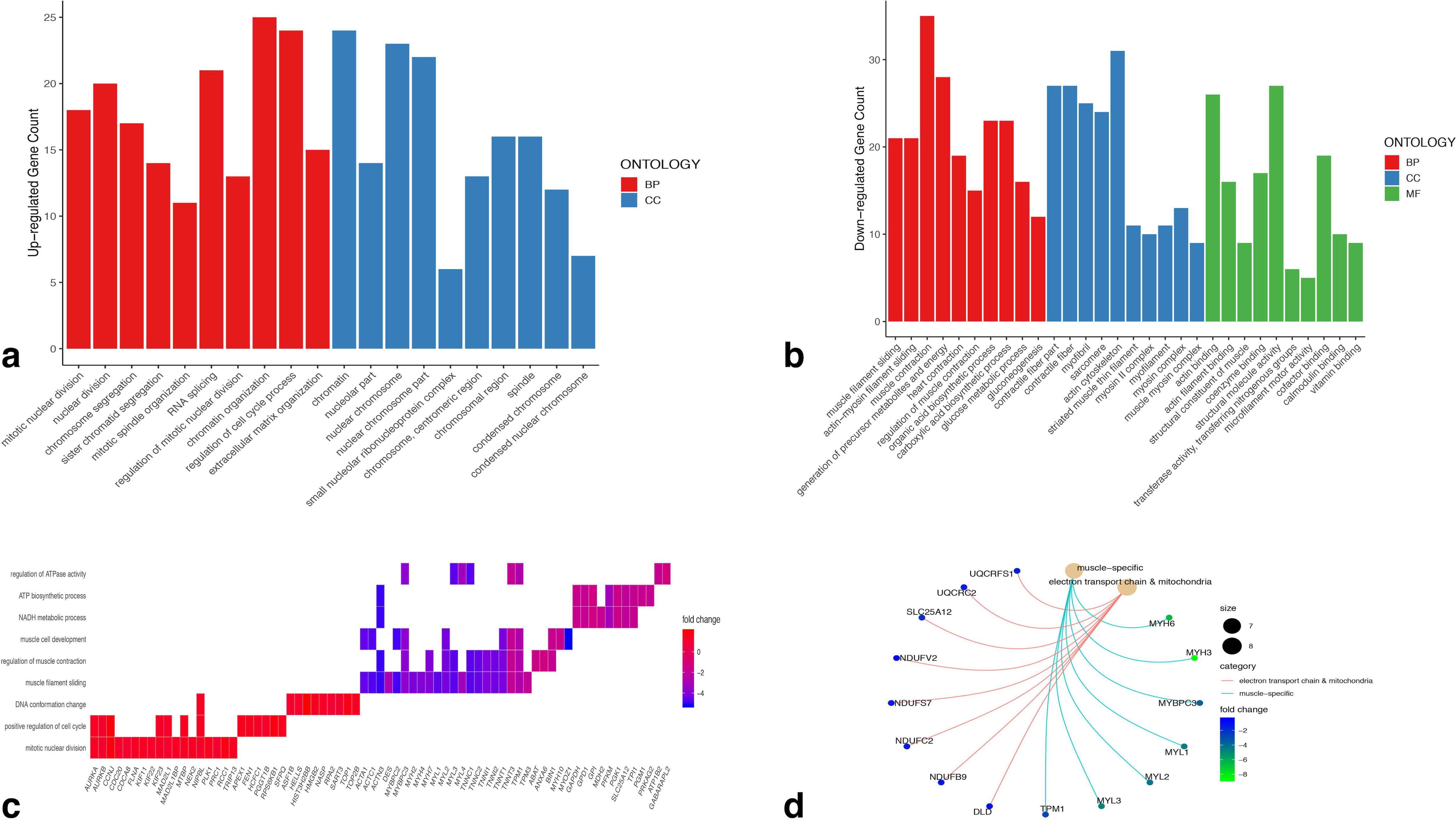
A) top 10 GO terms of each of biological processes and cellular components enriched by the top 181 up-regulated genes in regenerative vs. control comparison commonly identified by both technologies’ analyses. B) top 10 GO terms of each of biological processes, cellular components, and molecular functions enriched by the top 166 down-regulated genes in regenerative vs. control comparison commonly identified by both technologies’ analyses. C) Biological processes related to cell cycle, muscle-tissues, and metabolism enriched by some of the top up-regulated and down-regulated genes in regenerative vs. control comparison. D) some of the top DE muscle-specific genes and other DE metabolic genes being down-regulated in regenerative vs. control comparison at different rates.

The first principal component (PC1) as well as the sample-to-sample clustering heatmap of the z-scored, microarray DEA-based data show a clear separation among the samples based on their group types; for regenerative vs. control (**Figure 3A, B**) and for the other two comparisons (**Supplementary Figures 7A, B and 8A, B**). However, the second principal component (PC2) indicates no strong separation of any type. Therefore, differential gene expression between the group types is evidently the dominant source of variation in this case. Likewise, the separation among the samples displayed by the first principal component (PC1) in addition to the sample-to-sample clustering heatmap of the voom-counts, RNA-Seq DEA-based data, unlike the whole gene expression data-based approach, is apparently based on their group types, while the second principal component (PC2) seems to point towards no particular separation of any type; for regenerative vs. control (**Figure 3C, D**) and for the other two comparisons (**Supplementary >Figures 7C, D and 8C, D**). Thus, PC1 along with the heatmap clustering sufficiently and strongly demonstrate such high variation among the samples stems from differential expression between the group types.

**Figure 7:**
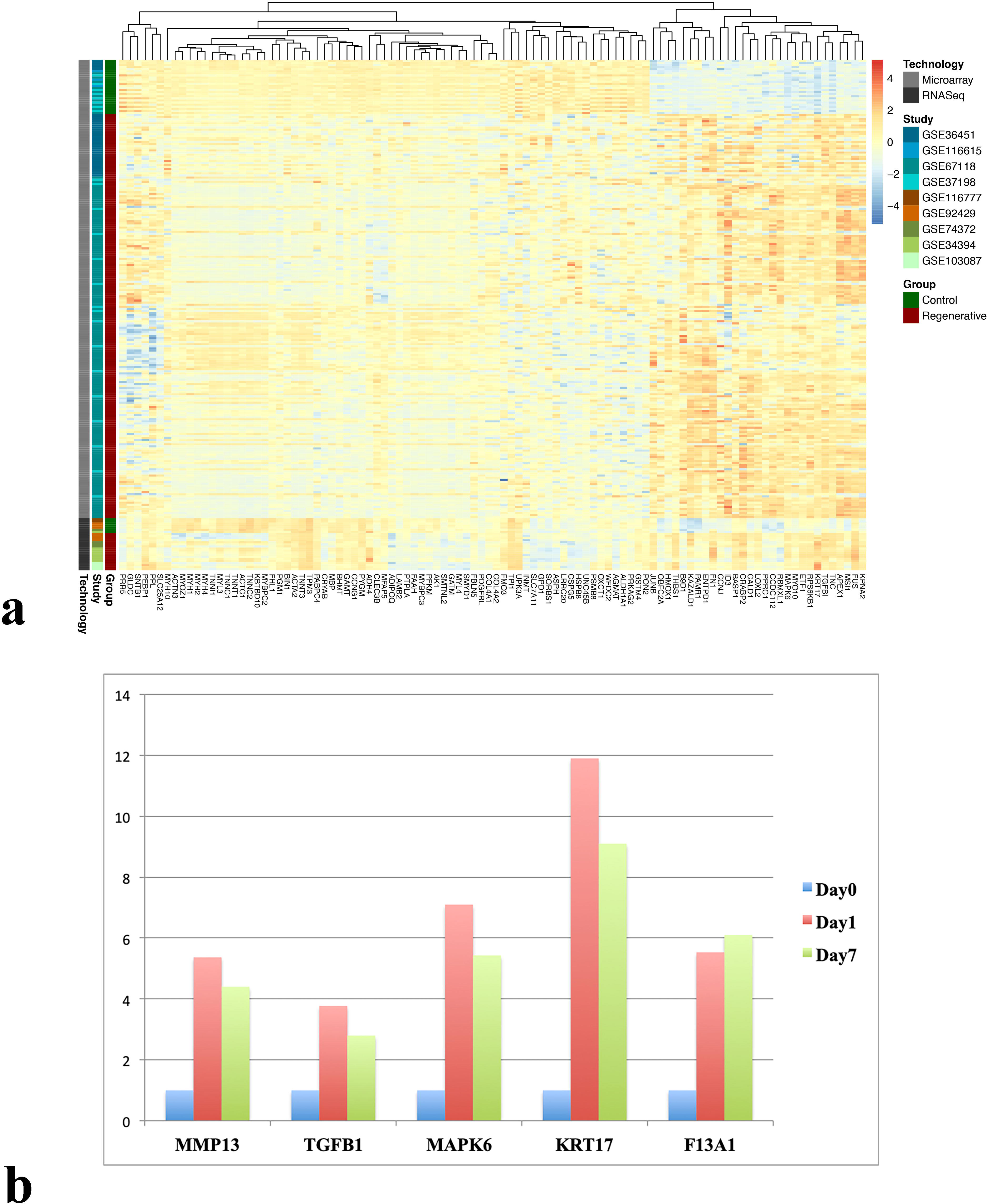
A) clustered heatmap of the top 100 DE genes commonly detected by the analyses of both technologies in regenerative vs. control comparison. B) qRT-PCR of selected 5 genes which are up-regulated at 1 and 7 dpa compared to day0.

Overall, the batch factor (study origin) has apparently been minimized during DEA as illustrated by the DEA-based PCA and clustering approach, which further demonstrated how group types are separated as a result of differential gene expression in both microarray and RNA-Seq data.

### 3.4 Distribution of DE genes is evident among all three comparisons

DE genes were found to be distributed among wound healing vs. control, regenerative vs. control, and regenerative vs. wound healing comparisons from both microarray (**Figure 4A**) and RNA-Seq (**Figure 4B**) data. This means that genes could possibly be differentially expressed in more than one comparison, simultaneously.

From microarray data, 42 genes were uniquely DE in wound healing vs. control, 105 genes were uniquely DE in regenerative vs. control, and 405 genes were uniquely DE in regenerative vs. wound healing. While regenerative vs. control and wound healing vs. control have 457 DE genes in common, regenerative vs. control and regenerative vs. wound healing have 1168 DE genes in common, and wound healing vs. control and regenerative vs. wound healing have 575 genes in common. All three comparisons shared 1018 DE genes.

From RNA-Seq data, 62 genes were uniquely DE in wound healing vs. control, 1226 genes were uniquely DE in regenerative vs. control, and 573 genes were uniquely DE in regenerative vs. wound healing. While 145 genes are commonly DE in regenerative vs. control and wound healing vs. control, 1582 genes are commonly DE in regenerative vs. control and regenerative vs. wound healing, and 177 genes are commonly DE in wound healing vs. control and regenerative vs. wound healing. All three comparisons shared 39 DE genes.

Although the distribution of DE genes in evident among comparisons, they may not necessarily share the same status of gene regulation; i.e., a gene can be up-regulated in one comparison and down-regulated in another one. In order to test this notion, correlations of the logFCs of the genes commonly DE in two or more comparisons were calculated in a pair-wise trend and visualized through scatter plots. Interestingly, the observed correlations within microarray data (**Figure 4C-H**) share a similar pattern of gene regulation directionality within RNA-Seq data (**Figure 4I-N**).

The Spearman correlation coefficients for logFCs of the genes commonly DE in regenerative vs. control and regenerative vs. wound healing in microarray and RNA-Seq data are 0.85 and 0.94, respectively (**Figure 4C,I**). Likewise, the coefficients for logFCs of the genes commonly DE in regenerative vs. control and wound healing vs. control in microarray and RNA-Seq data are 0.96 and 0.93, respectively (**Figure 4E,K**). This positive correlation indicates that genes which are DE in those comparisons mostly have the same direction of gene regulation (mostly up-regulated or down-regulated) with those in their respective comparison. On the other hand, the coefficients for logFCs of the 575 genes and 177 genes commonly DE in wound healing vs. control and regenerative vs. wound healing in microarray and RNA-Seq data are - 0.71 and - 0.95, respectively (**Figure 4D,J**). This negative correlation indicates that those genes are mostly oppositely regulated in these two different comparisons. The pair-wise correlations for the 1018 genes and 39 genes DE in all three comparisons in microarray (**Figure 4F-H**) and RNA-Seq (**Figure 4L-N**) data, respectively, demonstrate that those genes follow the same trend as when they are shared by only the two corresponding comparisons. From microarray data, the 1018 genes DE in regenerative vs. control and regenerative vs. wound healing are positively correlated (r_s_ = 0.52), in regenerative vs. control and wound healing vs. control are positively correlated (r_s_ = 0.77), and in regenerative vs. wound healing and wound healing vs. control are negatively correlated (r_s_ = - 0.02). From RNA-Seq data, the 39 genes DE in regenerative vs. control and regenerative vs. wound healing are positively correlated (r_s_ = 0.03), in regenerative vs. control and wound healing vs. control are positively correlated (r_s_ = 0.54), and in regenerative vs. wound healing and wound healing vs. control are negatively correlated (r_s_ = - 0.71).

### 3.5 Some DE Genes are Commonly Detected by the Analyses of Microarray and RNA-Seq data

Microarray and RNA-Seq DEA was carried out on the 4322 and 7562 genes, respectively. Since the same annotation source (AMBY_002a520748F probeset annotation file) was used for the analyses from both technologies, a total of 3653 genes are common between them (**Figure 5A**). Therefore, differing numbers of DE genes per comparison that would be commonly identified by the analyses of both technologies is conceivable. Indeed, after merging the DE gene list (adj P-value < 0.01) of microarray data with that of RNA-Seq data for each comparison, we found 170, 1254, and 1047 DE genes commonly identified by both technologies in wound healing vs. control, regenerative vs. control, and regenerative vs. wound healing, respectively (**Figure 5B-D**). Each set of those common DE genes per comparison were tested for their logFC correlation between both technologies and were visualized using scatter plots. The common DE genes of all three comparisons had positive Spearman correlations of their logFCs between the two technologies; r_s_ = 0.74 for wound healing vs. control, r_s_ = 0.71 for regenerative vs. control, and r_s_ = 0.77 for regenerative vs. wound healing (**Figure 5E-G**).

### 3.6 Identifying the Top DE Genes Commonly Detected by the Analyses of Microarray and RNA-Seq Data

In spite of having a significance level of an adjusted P < 0.01 of the DE genes detected by the analyses of both technologies for each comparison, many of those genes have low magnitudes of logarithmic fold changes (|logFC| < 1). Moreover, although the correlation of the logFCs of the DE genes between the two technologies is positive with a fairly high coefficient for each comparison, the number of genes with conflicting directionality of gene expression would still be an issue. Thus, in order to reliably call those genes “DE”, another criterion could be set through which the top significant genes can be extracted from the list of genes detected by both technologies, thereby, decreasing the number of genes with low logarithmic fold changes as well as those with opposite directions of gene expression. We, therefore, set the criterion as follows: first of all, genes with logFC magnitudes > 1 are separately extracted from both microarray and RNA-Seq DE lists. Next, the resultant two lists are merged to yield the top DE genes commonly detected by both microarray and RNA-Seq analyses. Based on this criterion, the extracted top DE genes commonly detected by both technologies are 91 genes in wound healing vs. control, 351 genes in regenerative vs. control, and 280 genes in regenerative vs. wound healing. Those genes along with their logFC from both technologies is listed in **Supplementary Table 6**. The correlation of the logFCs of those top DE genes between the two technologies is positive, with a Spearman’s coefficient of 0.44 for wound healing vs. control, 0.72 for regenerative vs. control, and 0.76 for regenerative vs. wound healing (**Figure 5H-J**). Those top DE genes were merged together to see whether they are distributed among the three comparisons. Indeed, some of those top DE genes are commonly DE in more than one comparison. However, notably, this time the number of top DE genes commonly DE in all three comparisons is zero, and the number of top DE genes specific to each comparison is relatively high (**Figure 5K**).

### 3.7 GO Terms are Enriched by the Top DE Genes of Each Comparison

Our top DE genes detected by the DEA of each of the two technologies enriched a variety of biological processes (BP), cellular components (CC), and molecular functions (MF), for each comparison (**Supplementary Tables 7-9**). Biological processes such as mitotic nuclear division, chromosome segregation, RNA splicing, regulation of cell cycle process, and extracellular matrix organization were found among the top 10 BPs enriched by the top 181 up-regulated genes (**Figure 6A**). Those up-regulated genes also enriched several cellular components, such as chromatin, nuclear chromosome, and spindle, all being among the top 10 CC categories (**Figure 6A**). On the other hand, muscle filament sliding, muscle contraction, heart contraction, generation of precursor metabolites and energy, glucose metabolic process, and gluconeogenesis were detected among the top 10 biological process enriched by the top 166 down-regulated regenerative vs. control genes (**Figure 6B**). Cellular components such as contractile fiber, actin cytoskeleton, and muscle-myosin complex were among the top 10 CC categories enriched by those downregulated genes (**Figure 6B**). The latter genes also enriched several molecular functions, including actin filament binding, microfilament motor activity, and calmodulin binding, all of which are among the top 10 MF categories (**Figure 6B**).

### 3.8 Heatmap of top 100 and qPCR results

In order to look at the most differentially expressed genes detected by the DEA of both microarray and RNA-Seq data in regenerative samples compared to the controls all in a single and interpretable plot, the top 100 DE genes were selected and visualized in a gene-wise hierarchically-clustered heatmap (**Figure 7A**). The clustering shows an overall conspicuous clustering between regenerative and control samples from data of each technology. qRT-PCR of selected genes at 0,1, and 7 dpa was performed to validate the IDA approach. MMP13, TGFB1, MAPK6, Keratin17 and Coagulation factorXIII were used to test the accuracy of our analysis. Expression levels of all these genes were upregulated at day1 and day7 compared to day0 (**Figure 7B**), which is consistent with output of our analyses.

## 4. DISCUSSION

This study reports an integrative data analysis on axolotl gene-expression data. To our knowledge, this is the first study describing an integrative analysis methodology by which publicly available microarray and RNA-Seq axolotl data were leveraged in order to identify differentially expressed genes marking the wound healing and regenerative phases of the axolotl limb.

Despite the more frequent and persistent application of late-stage data integration (aka “meta-analysis”) in the literature in contrast to early-stage data integration (aka “merging”) [40], the latter has been applied in several studies [40,75–77] and was chosen for the purpose of our study. Previous studies have propounded that Late-stage data integration may perform less competently than early-stage data integration when the goal of the analysis is to discover robust biomarkers [40]. It has been postulated that computing separate statistics and taking the average is often less powerful compared to aggregation of data as a first step and then deriving statistics from this data [40,78]. For instance, a higher number of differentially expressed genes was detected by early-stage integrative data analysis as reported by Taminau *et al* [39]. Another advantage of the merging method is facilitating the application of prediction models which were originally built for a subset of studies onto studies from different platforms [40,42].

While the merging method recognizes all samples coming from different datasets across different platforms as a single dataset when testing the same hypothesis, the existence of systematic biases may introduce unwanted batch effects (non-biological differences) during the analysis of gene signatures and, consequently, true biological differences can be masked among the conditions of interest [39,40,48,79]. Moreover, several intra-laboratory variables that may have ambiguous or absent GEO entries such as amputation site, size, feeding, and maintenance protocols of axolotls are amongst many factors which can influence how control and test samples cluster together, and probably contributing to the observed batch effect in whole gene expression data-based PCA and clustering heatmap (**Figure 2 and Supplementary Figures 3 and 4**). Nonetheless, preservation of true biological (gene-expression) differences between experimental groups were successfully attained after DEA and visualized through PCA and clustering heatmap (**Figure 3 and Supplementary Figures 7 and 8**), indicating an indecisive role of any intra-laboratory variables. Notably, minimization of batch effects and maximization of true gene-expression differences were mainly due to the application of transformation and normalization techniques during data processing [40,64] along with accounting for the experiment source (study origin) for each sample while performing DEA [69].

The consistency of the results obtained from each of microarray and RNA-Seq DEAs with one another indicates a fairly valid approach of integrative analysis that we took. First and foremost, the correlations of each pair-wise distribution of the DE genes among all three comparisons from microarray data (**Figure 4A, C-H**) accord with those from RNA-Seq data (**Figure 4B, I-N**). Positive correlations are always observed among the genes commonly DE in regenerative vs. control and regenerative vs. wound healing (**Figure 4C, F, I, L**), as well as among those commonly DE in regenerative vs. control and wound healing vs. control (**Figure 4E, G, K, M**). Further, negative correlations are always observed among the genes commonly DE in regenerative vs. wound healing and wound healing vs. control (**Figure 4D, H, J, N**). This also suggests a possibly true underlying biological behavior of those genes in their respective comparisons. Secondly, DE genes commonly detected by DEA of each of the two technologies are always positively correlated between microarray and RNA-Seq data, along with their top DE genes, for every comparison (**Figure 5B-J**). Several previous studies have also reported such strong positive correlations between microarray and RNA-Seq data [80–83]. Although most of the DE genes are positively correlated between the two technologies for each comparison, small number of genes have opposite gene expression direction, which is probably attributable to artifacts of gene expression technologies. Nevertheless, the number of those genes with an opposite direction was found significantly low for the top DE genes (**Figure 5H-J**). Even more, the distribution of those top DE genes among all three comparisons suggests a pattern in which the more top DE genes are extracted the more specialized those genes become to a particular comparison (**Figure 5K**).

Some of the top DE genes detected by both technologies in wound healing vs. control comparison concur with previously identified genes in the wound healing response. In wound healing process, at around 6-8 hours post-amputation (hpa), epithelial cells tend to migrate to the amputation site to form the “wound epithelium (WE)” beneath which are cellular and extracellular debris as well as a damaged vasculature [54,84]. Basel cells of this wound epidermis lacking a basement membrane at 24 hpa are characterized with highly up-regulated thbs1 gene [53,85], which was found as a top up-regulated gene in our wound healing vs. control list. Furthermore, matrix metalloproteinases activity is required to regulate the extracellular matrix and these proteins are enriched in the wound epidermis as soon as an injury takes place [86]. Our wound healing vs. control top DE genes list also included metalloproteinases such as mmp1, mmp3, mmp19, and their regulator timp1, in addition to the extracellular matrix-generating component tnc [53]. Complete list of the top DE genes in wound healing vs. control is presented in **Supplementary Table 6** and GO terms enriched by them is documented in **Supplementary Table 7**.

Genes which were previously implicated in the regenerative process also concord with some of our top DE genes detected by both technologies in regenerative vs. control comparison. Following two days post-amputation, processes such as DNA replication, mitosis, and cell cycle are enriched by a set of up-regulated genes the majority of which signify a transition phase in the limb regeneration program during 2-3 dpa interval [16]. After 3 dpa, those genes either undergo an increased rate of up-regulation or sustain a relatively constant expression until 28 dpa [16]. Concordantly, many gene ontology terms, particularly those related to cell cycle, were enriched by many of our top up-regulated regenerative vs. control genes, such as mitotic nuclear division, positive regulation of cell cycle, and DNA conformation change (**Figure 6C**). This punctuated increase of cell cycle transcripts is, therefore, indicative of a striking change in the population of proliferative cells taking place at the stump of the distal limb [16]. Besides, earlier studies have reported significant reduction of muscle-specific genes over the course of limb regeneration [15,16,53,57,87]. During early response to limb amputation, the limb stump also undergoes muscle tissue remodeling along with diminished levels of muscle-specific transcripts [53]. The reduction in the expression of muscle-specific transcripts, however, becomes so significant by around 10 dpa which strongly implies a complete absence of muscle tissues [16]. Therefore, depletion of muscle tissues in such an absolute manner is propounded to be an essential step towards the recruitment of progenitor cells and the initiation of blastemal outgrowth [53]. This observation is also in line with our top down-regulated regenerative vs. control genes which enriched many muscle-specific GO terms, including muscle filament sliding, regulation of muscle contraction, and muscle cell development (**Figure 6C**). In addition to the reduction of muscle-specific transcripts by 10 dpa, expression levels of some transcripts associated with metabolic processes also markedly drop by around the same time [16,53]. Indeed, some of our top down-regulated regenerative vs. control genes enriched several GO terms associated with cellular metabolic processes, such as NADH metabolic process, ATP biosynthetic process, and regulation of ATPase activity (**Figure 6C**).

Another interesting observation is that cellular metabolic genes are still expressed by the cells which do not die or undergo reprogramming. Therefore, non-absolute, moderate depletion of some metabolic genes in contrast to the complete reduction of muscle-specific genes during the regenerative phase was previously shown [53]. This relationship was also discovered in our regenerative vs. control comparison between top down-regulated muscle-specific genes and some of the metabolic genes functioning in electron transport chain and mitochondria, most of which are not among the top DE genes (**Figure 6D**). As depicted, those metabolic genes are much less down-regulated than the muscle-specific genes during the regenerative phase.

When discussing “top DE genes” or “DE genes”, we refer to those genes commonly detected by the DEA of each of microarray and RNA-Seq. However, an important point to underline here is that two of the muscle-specific genes (myh6, myh3) and a metabolic gene (ndufv2) from **Figure 6D** were detected DE (adjusted P < 0.01) only by RNA-Seq DEA. Since Affymetrix and Agilent data’s gene lists were merged together, the resultant microarray genes did not completely match those of RNA-Seq which were annotated using the Affymetrix annotation file. Consequently, many genes which were used for RNA-Seq DEA were not represented in the final 4322 microarray gene list. This is, therefore, indicative of the fact that the reliability of our results may be extended to those DE genes specifically detected by only RNA-Seq DEA, and not to be limited to those commonly detected by the DEA of each of the two technologies. Excluding the Agilent dataset from microarray DEA could -probably-lead to less complicated downstream analysis. On the other hand, sample size is often a key for and a fundamental premise of having a successful and powerful integrative analysis; and in this study, this concept was further fulfilled by integrating the Agilent dataset.

There are some inherent limitations about this study that need to be addressed. Despite our successful minimization, the merging methodology cannot guarantee the complete removal of between-laboratory heterogeneity (batch effects) across experiments. In some cases, the quality of the deposited, original data could play a deterministic role in the acquisition of the most statistically-sound downstream results. The major limitation to underline, however, lies within the concept of data integration itself along with the gene-level data processing approach we took. Integrative analysis of different platforms provides statistical power through increased sample size at the expense of having a large number of genes for differential expression and downstream analyses. The process of merging the genes of Affymetrix platform with those of Agilent platform showed a decrease in the total number of microarray genes which were used for DEA. Moreover, the approach of annotating our integrated RNA-Seq data from the 31k contigs-V3 probesets alignment eliminated a plethora of transcripts which could be important in the limb regeneration process. Therefore, a “meta-analysis” approach which would utilize probeset-level information for separately processing data from each platform, from each technology, and then combining the resultant statistics could solve the aforementioned issues. That being said, since we rather chose the “merging” approach on the premise that it is well suited for biomarker discovery, collapsing information from prove-level to gene-level was an absolute necessity due to platform design variations and for direct comparability with RNA-Seq DEA results. It is also plausible to conceive the RNA-Seq annotation process from the list of Affymetrix-Agilent genes instead of the whole Affymetrix annotation file, which would also lead to a decrease in the total number RNA-Seq genes to be used for DEA, not to mention the further decrease that normally comes from the filtering steps. Therefore, in this study, we tried to sustain a trade-off between increased statistical power through integration of data and having the maximum possible number of genes to be analyzed.

## CONCLUSION

To the best of our current knowledge, this is the first study describing an integrative analysis of publicly available microarray and RNA-Seq axolotl data, with an aim to uncover differentially expressed genes signifying the wound healing and regenerative phases of the axolotl limb. The validity of our DEA methodology can be realized from observing the same distribution pattern of DE genes within the DEA of each technology and the positive correlation of the DE genes along with the top ones between both technologies for all three comparisons; wound healing vs. control, regenerative vs. control, and regenerative vs. wound healing. Our results included many DE genes enriched in a wide variety of biological processes in accord with those described in previous studies. From such concordance, we also suggest considering the results of RNA-Seq DEA to be as reliable as those commonly detected by DEA of both technologies. qPCR experiment (**Figure 7B**) provides another layer of evidence to validate our computational findings. Future direction of this study would aim to functionally validate more of our top DE genes to possibly identify some of them as biomarkers for the regenerative phases of the axolotl limb. Besides, other statistical approaches can be applied as an extension to the described methodology, such as using ANOVA test to detect DE genes among all experimental groups at once that could provide an enhanced, unified interpretation and visualization of the data. We hope for this study to provide the required confidence for researchers around the world to consider our DE lists for their functional studies. We also think that researchers can integrate their data into our methodology, thereby increasing sample size for more statistical confidence and better biomarker discovery.

**Supplementary Figure 1:** Detailed workflow for Affymetrix and Agilent Microarray axolotl data processing prior to differential expression analysis.

**Supplementary Figure 2:** Detailed workflow for Illumina RNA-Seq data processing prior to differential expression analysis.

**Supplementary Figure 3:** Whole gene expression data-based principal component analysis and sample-to-sample clustering heatmaps for control and wound healing samples of A,B) Microarray quantile-normalized, log2-transformed, z-scored data (4322 genes, 119 samples) and of C,D) RNA-seq logCPM (voom) counts data (7562 genes, 14 samples). A,C) principle component analysis. B,D) sample-to-sample clustering heatmap. The number of samples per group are; 26 Control and 93 wound healing for microarray data; 7 control and 7 wound healing for RNA-Seq data.

**Supplementary Figure 4:** Whole gene expression data-based principal component analysis and sample-to-sample clustering heatmaps for regenerative and wound healing samples of A,B) Microarray quantile-normalized, log2-transformed, z-scored data (4322 genes, 287 samples) and of C,D) RNA-seq logCPM (voom) counts data (7562 genes, 25). A,C) principle component analysis. B,D) sample-to-sample clustering heatmap. The number of samples per group are; 194 regenerative and 93 wound healing for microarray data; 18 regenerative and 7 wound healing for RNA-Seq data.

**Supplementary Figure 5:** Raw P and adjusted-P value distribution of microarray DEA results. p-value histogram (left) and adj.P.Value histogram (right) were plotted after DEA on Microarray data for the following comparisons A,B) wound healing vs. control, C,D) regenerative vs. control, E,F) regenerative vs. wound healing.

**Supplementary Figure 6:** Raw P and adjusted-P value distribution of RNA-Seq DEA results. p-value histogram (left) and adj.P.Value histogram (right) were plotted after DEA on RNA-Seq data for the following comparisons A,B) wound healing vs. control, C,D) regenerative vs. control, E,F) regenerative vs. wound healing.

**Supplementary Figure 7:** DEA-based principal component analysis and sample-to-sample clustering heatmaps for wound healing vs. control comparison of A,B) Microarray quantile-normalized, log2-transformed, z-scored data after DEA (2092 DE genes, 119 samples) and of C,D) RNA-seq logCPM (voom) counts data after DEA (423 DE genes, 14 samples). A,C) principle component analysis. B,D) sample-to-sample clustering heatmap. The number of samples per group are; 26 control and 93 wound healing for microarray data; 7 control and 7 wound healing for RNA-Seq data. The DE genes have an adjusted p-value < 0.01.

**Supplementary Figure 8:** DEA-based principal component analysis and sample-to-sample clustering heatmaps for wound healing vs. control comparison of A,B) Microarray quantile-normalized, log2-transformed, z-scored data after DEA (3166 DE genes, 287 samples) and of C,D) RNA-seq logCPM (voom) counts data after DEA (2371 DE genes, 25 samples). A,C) principle component analysis. B,D) sample-to-sample clustering heatmap. The number of samples per group are; 194 regenerative and 93 wound healing for microarray data; 18 regenerative and 7 wound healing for RNA-Seq data. The DE genes have an adjusted p-value < 0.01.

